# Inhibition of NLRP3 Differentially Regulates Blood Pressure and Inflammation in Male versus Female DOCA-Salt Sprague Dawley Rats

**DOI:** 10.64898/2026.03.17.712521

**Authors:** Cameron M. Liss, Ahmed Elmarakby, Kelly Sullivan, Peyton Hanson, Kasey Belanger, Riyaz Mohamed, David L. Mattson, Erin B. Taylor, Michael J. Ryan, Jennifer C. Sullivan

**Affiliations:** Department of Physiology, Medical College of Georgia at Augusta University, Augusta, Georgia; Department of Oral Biology, Dental College of Georgia at Augusta University, Augusta, Georgia; Department of Physiology and Biophysics, University of Mississippi Medical Center, Jackson, Mississippi; Columbia VA Health Care System, Columbia, South Carolina; University of South Carolina School of Medicine, Columbia, South Carolina

**Keywords:** Hypertension, MCC950, kidney, T cells, sex

## Abstract

**Background:** Deoxycorticosterone acetate (DOCA)-salt induces greater increases in blood pressure (BP) and a more pro-inflammatory T cell profile in males compared to females. T cells contribute to DOCA-salt hypertension, however, the mechanisms driving T cell activation remain unclear. The NLRP3 inflammasome has been implicated in DOCA hypertension in male mice. Little is known regarding NLRP3 in females. The goal of the current study was to test the hypothesis that NLRP3 contributes to greater increases in BP and renal inflammation with DOCA in males vs. females.

**Methods:** Renal NLRP3 protein levels were measured in normotensive and hypertensive male and female subjects and in male and female Sprague Dawley uni-nephrectomized (UNX) control and DOCA-salt rats. Additional 11-wk-old Sprague Dawley rats were UNX and randomized to: 1) DOCA + vehicle or 2) DOCA + the NLRP3 inhibitor MCC950 (10 mg/kg/day in saline) from 11-14 wks of age. At 14-wks-of-age rats were euthanized, terminal plasma samples and remaining kidneys were collected for flow cytometric analysis of T cells.

**Results:** Renal NLRP3 levels were significantly greater in hypertensive males and females vs. normotensive controls. DOCA increased BP in both sexes, with greater elevations in males. MCC950 attenuated DOCA-induced increases in BP in male, but not female rats. MCC950 decreased circulating and renal CD4⁺ and Th17 cells in both sexes, although the effect was greater in males.

**Conclusion:** Despite both males and females exhibiting an increase in NLRP3 in hypertension, NLRP3 contributes to BP elevations only in DOCA-salt males.

## Introduction

Sex differences in blood pressure (BP) regulation are well established, with both clinical and experimental studies consistently reporting higher BP in young males compared with age-matched females ^1–3^. Deoxycorticosterone acetate (DOCA)-salt, a low renin model of hypertension, is a widely used experimental model to study mineralocorticoid-driven hypertension, inflammation, and renal injury ^4^. Moreover, males exhibit greater increases in BP in DOCA-salt hypertension compared to females ^1,5^, yet the mechanisms responsible for this sex difference remain unclear.

T cells play a central role in the development and progression of DOCA-salt hypertension ^1,6^ and T cells have been shown to contribute to sex differences in hypertension ^1,7^. Indeed, we have previously shown that female DOCA-salt hypertensive rats have more anti-inflammatory T regulatory cells (Tregs) than males, and decreasing Tregs results in an increase in BP only in females abolishing the sex difference in BP ^1^. Yet, what drives the sex difference in the T cell profile has not been investigated.

The NLRP3 inflammasome is a well-characterized cytosolic sensor, expressed primarily in innate immune cells that detects and responds to pathogen- and danger-associated molecular patterns (PAMPs and DAMPs) released from stressed or damaged cells ^8^. NLRP3 activation leads to the production of IL-1β and IL-18 to propagate an inflammatory response ^9,10^. NLRP3 activation has been implicated in the development and progression of numerous chronic low grade inflammatory diseases ^9–13^. Importantly, NLRP3 has been shown to contribute to DOCA-salt-induced increases in BP and renal inflammation and injury in male mice ^9,10^.

While studies show that NLRP3 contributes to sex differences in nitroglycerine-induced migraines ^14^ and in atherogenesis in mice ^15^, a role for NLRP3 in sex differences in BP has not been explored. Therefore, the goals of the current study were to determine if NLRP3 expression increases in hypertension and to assess the relative contribution of NLRP3 to DOCA-salt induced increases in BP and the renal T cell profile in male and female rats. We hypothesize that greater NLRP3 in males leads to greater increases in BP and a more pro-inflammatory T cell profile with DOCA-salt hypertension vs. females.

## Methods

This study was conducted in compliance with ethical guidelines and approved by the Augusta University Institutional Animal Care and Use Committee (IACUC). Initially, 11-week-old male and female Sprague-Dawley (SD) rats (Envigo, Indianapolis, IN) underwent uni-nephrectomy (UNX) surgery and were randomized to receive either tap water to drink and serve as controls or a 21-day slow-release DOCA pellet (200 mg, S.C.) with saline to drink (n =5/group). After 3 weeks, rats were euthanized and the remaining kidneys were harvested to measure renal NLRP3 using a commercial rat ELISA (LSBIO, MA; Cat # LS-F39627). Kidney samples were homogenized in PBS and results are reported as pg/mg protein.

NLRP3 levels were also measured in human kidneys obtained from LifeLink® Foundation, which provides non-transplantable human donor organs for research purposes. LifeLink® is a Centers of Medicare & Medicaid Services certified Organ Procurement Organization for the state of Georgia whose primary mission is to recover donor organs and tissues for human transplantation. Upon authorization for research, human kidneys deemed unsuitable for transplant with corresponding-deidentified clinical and medical history were made available to Augusta University by LifeLink®, an approved research entity that has agreed to receive such donor organs and tissues for use in studies under Augusta University Institutional Review Board # 1732545. Twenty-two kidney samples from normotensive (BP ≤ 120/80 mm Hg) and hypertensive (BP≥ 140/90 mm Hg) male and female subjects were obtained and renal NLRP3 levels were measured via human ELISA (LSBIO, MA; Cat # LS-F31954). Samples were homogenized in PBS and results are reported as ng/mg protein.

Additional 11-week-old male and female SD rats underwent UNX and were randomized to two treatment groups (n =6/group): (1) DOCA-salt with saline to drink or (2) DOCA-salt plus the NLRP3 inhibitor MCC950 (10 mg/kg/day in saline). This dose was selected based on previous publications ^9,10^. An additional group of rats was implanted with radio-telemetry transmitters for continuous BP monitoring as previously described ^1^. Water intake was measured daily, and body weight was recorded every three days to maintain consistent MCC950 dosing. After three weeks of treatment, all rats were euthanized under isoflurane anesthesia. A thoracotomy was performed, and a terminal blood sample was collected. The remaining kidney was harvested and processed for flow cytometric analysis of T cells, biochemical assessments, and histopathological analysis of renal structure.

### Biochemical Analyses

Kidney slices were homogenized in cold PBS and NLRP3 (LSBIO, MA, Cat # LS-F39627), IL-1β (Thermo-Fisher, MA, Cat # BMS630) and IL-18 (Abcam, MA, Cat # 214028) protein levels were measured by ELISA and reported as pg/mg or ng/mg protein. Caspase-1 activity in renal homogenates was analyzed with the Caspase-Glo 1 inflammasome assay (Promega, WI, Cat # G9951) according to the manufacturer’s protocol and results are reported as relative fluorescence units/µg protein.

### Analytical Flow Cytometry

Terminal blood and kidney samples were collected for flow cytometric assessment of the T cell profile. Single-cell suspensions were generated from kidneys by mechanical dissociation, filtration through a 100 μm cell strainer (BD Biosciences, San Diego, CA), and centrifugation as previously described ^1,16^. To assess T cell phenotype, cells were incubated with antibodies targeting CD3 (1:100, eBioscience, San Diego, CA) and CD4 (1:100, BD Biosciences, San Diego, CA). After washing, cells were fixed and permeabilized using fix/perm concentrate (eBioscience, San Diego, CA) before staining for intracellular markers: Foxp3 (1:100, eBioscience, San Diego, CA) to identify Tregs (expressed as %CD3+CD4+) and RORγt (1:100, eBioscience, San Diego, CA) to identify T helper (Th)17 cells. After staining, cells were washed and analyzed using a flow cytometer (FACS Calibur, BD Biosciences, San Jose, CA), with data collected using Cell Quest software. Flow cytometry data were processed and analyzed using Flow-Jo 10.5.3 software (BD Company, Franklin Lakes, NJ).

### Assessment of Renal Injury

A portion of the remaining kidney was fixed in 10% buffered formalin overnight and embedded in paraffin wax. Paraffin-embedded kidney sections (10 μm thick) were stained with Periodic Acid-Schiff Hematoxylin (PASH) to assess glomerular injury. Approximately ten images per section were captured at 200X magnification, and glomeruli within each image were blindly scored on a 1–4 scale, where 1 = normal, 2 = mild mesangial expansion, 3 = segmental sclerosis, and 4 = global sclerosis.

To evaluate renal fibrosis, additional 10 μm sections were stained with Masson trichrome to assess collagen deposition. Ten images per slide were collected at 200X magnification, and the degree of fibrosis was blindly scored on a 1–5 scale, where 1 = minimal collagen deposition and 5 = extensive interstitial fibrosis with dense blue staining.

We measured plasma and urinary creatinine using QuantiChrom^TM^ Creatinine assay kit (Bioassay System, CA, Cat # DICT500) and values were used to calculate creatinine clearance. Urinary protein concentrations were also measured using Pierce BCA Protein Assay (Thermo-Fisher, MA, Cat # 23223).

### Statistical Analysis

Telemetry data were analyzed using repeated-measures analysis of variance (ANOVA). Flow cytometry, ELISA, and histological outcomes were analyzed using two-way ANOVA to assess main effects of sex and treatment, followed by Tukey’s post hoc multiple comparisons test. Analyses were performed using GraphPad Prism Version 10 software (GraphPad Software Inc, La Jolla, CA). All data are expressed as mean ± SEM, and differences were considered statistically significant when p<0.05.

## Results

### Renal NLRP3 is greater in hypertensive subjects and DOCA-salt rats vs. normotensive controls

Renal NLRP3 levels were measured by ELISA in kidneys from hypertensive and normotensive men and women and male and female SD rats. NLRP3 levels were significantly greater in hypertensive subjects vs. normotensive controls, with comparable increases in men and women (Figure 1 A, P_BP_= 0.0005, P_sex_= 0.78, P_interaction_= 0.50). Similarly, renal NLRP3 levels were greater in male and female DOCA-salt hypertensive rats vs. control UNX rats with no differences between males and females (Figure 1B, P_BP_< 0.0001, P_sex_= 0.51, P_interaction_= 0.85).

**Figure 1:**
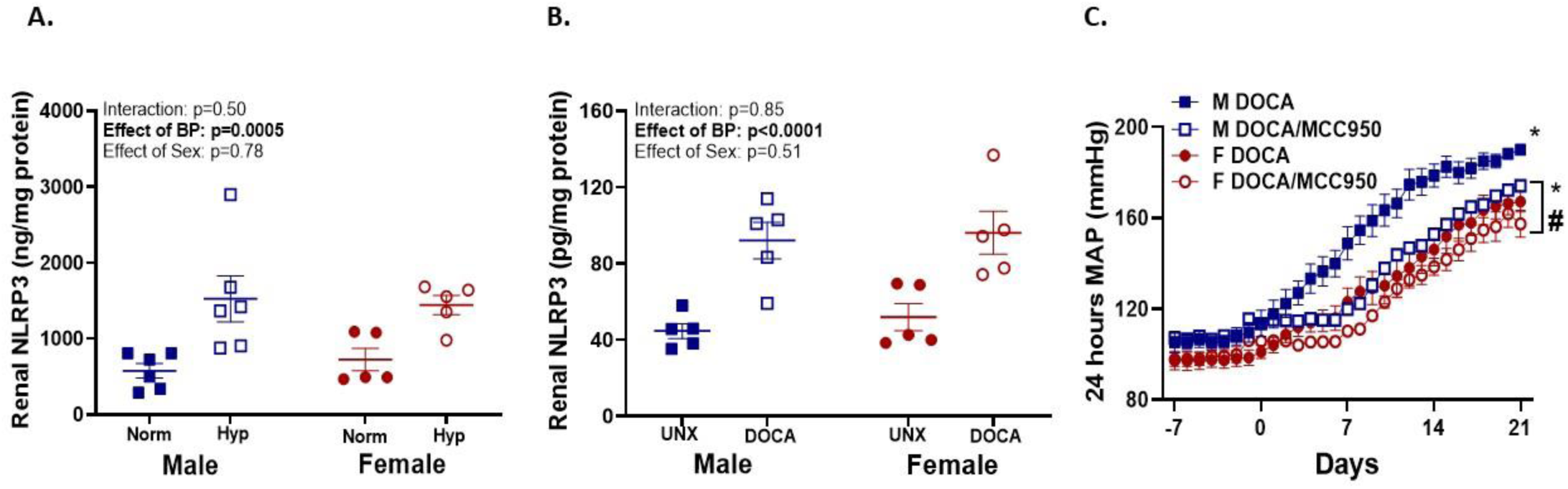
Renal NLRP3 increases in hypertension and pharmacological inhibition of NLRP3 attenuates DOCA-salt-induced increases in BP only in male rats. Renal NLRP3 protein levels were measured by ELISA in male and female normotensive (Norm) and hypertensive (Hyp) subjects (A, n = 5-6/group) and uni-nephrectomized (UNX) male (M) and female (F) Sprague Dawley (SD) rats following 3 weeks of control or deoxycorticosterone acetate (DOCA; 200 mg, S.C.) treatment (B, n = 5/group). Data are expressed as mean ± SEM and analyzed using 2-way ANOVA. Mean arterial pressure (MAP) was measured via telemetry (C) in DOCA treated M and F SD DOCA-salt rats randomized to saline control or the NLRP3 inflammasome inhibitor MCC950 (10 mg/kg/day in saline) for 3 weeks (n = 5/group). Data are expressed as mean ± SEM and analyzed using ANOVA for repeated measures. For all studies, P < 0.05 is considered significant. * indicates P < 0.05 vs. corresponding baseline values. ^#^ indicates P < 0.05 vs. male rats.

### Inhibition of NLRP3 attenuates DOCA-salt-induced increases in BP in male, but not female rats

To determine the contribution of NLRP3 to DOCA-salt-induced increases in BP, additional male and female DOCA-salt rats were randomized to treatment with vehicle or the NLRP3 inhibitor, MCC950. Three weeks of DOCA-salt treatment increased BP in both male and female rats compared to baseline values (mm Hg: 190.1±1.2 vs. 113.6±5.8, P< 0.0001 in males and 167.3±4.5 vs. 101.1±2.9, P< 0.0001 in females), with males exhibiting greater increases in BP with DOCA treatment than females (P_sex_< 0.01, Figure 1C). Despite comparable increases in NLRP3 in male and female rats following 3 weeks of DOCA-salt treatment, MCC950 attenuated DOCA-salt-induced increases in BP in male (mm Hg: 174.3±1.6 vs. 190.1±1.2), but not female rats (mm Hg: 157.5±5.9 vs. 167.3±4.5) abolishing sex differences in BP (P_treatment_> 0.003, P_interaction_= 0.04, Figure 1C).

NLRP3 activation drives the activation of caspase-1, leading to the production of IL-1β and IL-18 ^9,10^. To confirm the effectiveness of MCC950, renal NLRP3, IL-1β, and IL-18 expression and caspase activity were measured at the end of the treatment period in vehicle- and MCC950-treated DOCA-salt rats. Although MCC950 attenuated DOCA-induced increases in BP only in males, MCC950 resulted in similar decreases in NLRP3 protein levels in male and female DOCA-salt rats (P_treatment_< 0.0001; P_sex_= 0.98; P_interaction_= 0.81, Figure 2A). Consistent with protein data, MCC950 resulted in similar decreases in NLRP3 mRNA expression levels in male and female DOCA-salt rats (Figure S1A). MCC950 attenuated DOCA-salt-induced increases in IL-1β protein (P_treatment_= 0.0009; P_interaction_= 0.83, Figure 2B) and mRNA expression (Figure S1B), IL-18 (P_treatment_< 0.0001; P_interaction_= 0.13, Figure 2C), and caspase-1 activity in both sexes (P_treatment_= 0.0018, P_interaction_= 0.65, Figure 2D). IL-1β protein levels and caspase activity were greater in males vs. females regardless of treatment (P_sex_= 0.006 and P_sex_= 0.05, respectively), whereas IL-18 protein levels were higher in females (P_sex_= 0.045).

**Figure 2:**
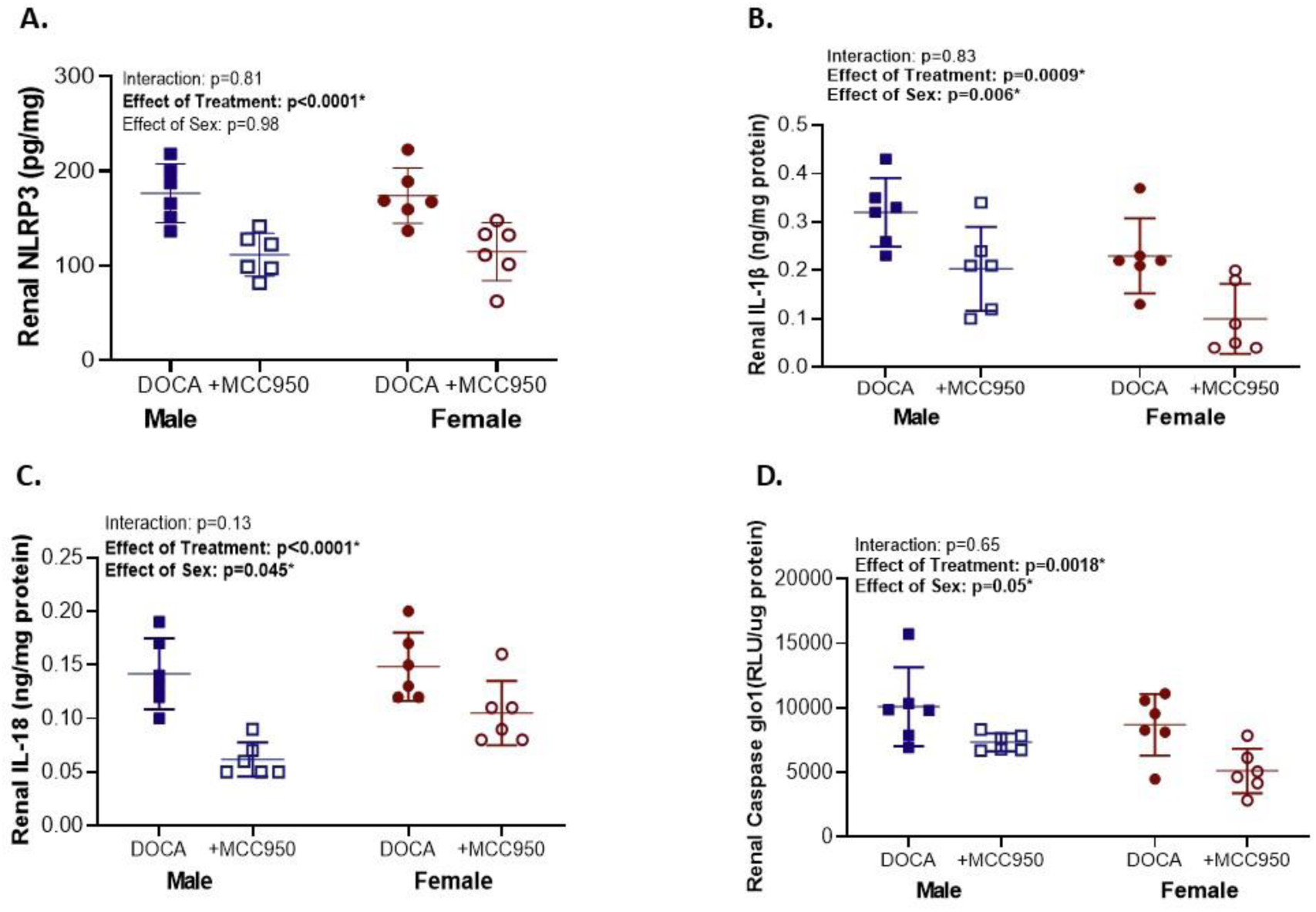
Oral administration of MCC950 (10 mg/kg/day) inhibits the activation of the NLRP3 inflammasome in male and female DOCA-salt hypertensive rats. Renal NLRP3 (A), IL-1β (B) and IL-18 (C) protein levels were measured by ELISA and renal caspase-1 activity was measured by caspase-Glo 1 inflammasome assay (D) in kidneys from deoxycorticosterone acetate (DOCA) treated male and female Sprague-Dawley (SD) rats randomized to saline control or the NLRP3 inflammasome inhibitor MCC950 (10 mg/kg/day in saline) for 3 weeks (n = 6). Data are expressed as mean ± SEM and analyzed using 2-way ANOVA. * indicates significance.

### MCC950 reduces circulating pro-inflammatory T cells in DOCA-salt hypertensive males and females

Circulating T cells were measured by flow cytometry in vehicle- and MCC950-treated DOCA-salt rats. MCC950-treated rats had significantly fewer CD3^+^ (P_treatment_= 0.004), CD4^+^ (P_treatment_< 0.0001), and Th17 cells (P_treatment_< 0.0001) in whole blood vs. vehicle control treated DOCA-salt rats (Figure 3A-C). Females had a greater percentage of CD3^+^ T cells than males (P_sex_=0.005), although the effect of MCC950 on CD3^+^ (P_interaction_= 0.72), CD4^+^ (P_interaction_= 0.32) and Th17 cells (P_interaction_= 0.13) were comparable between the sexes (Figure 3A-C). Circulating Tregs were not altered by MCC950 in either sex (P_treatment_= 0.84) and percentages were comparable between males and females (P_sex_= 0.31, P_interaction_= 0.67, Figure 3D).

**Figure 3:**
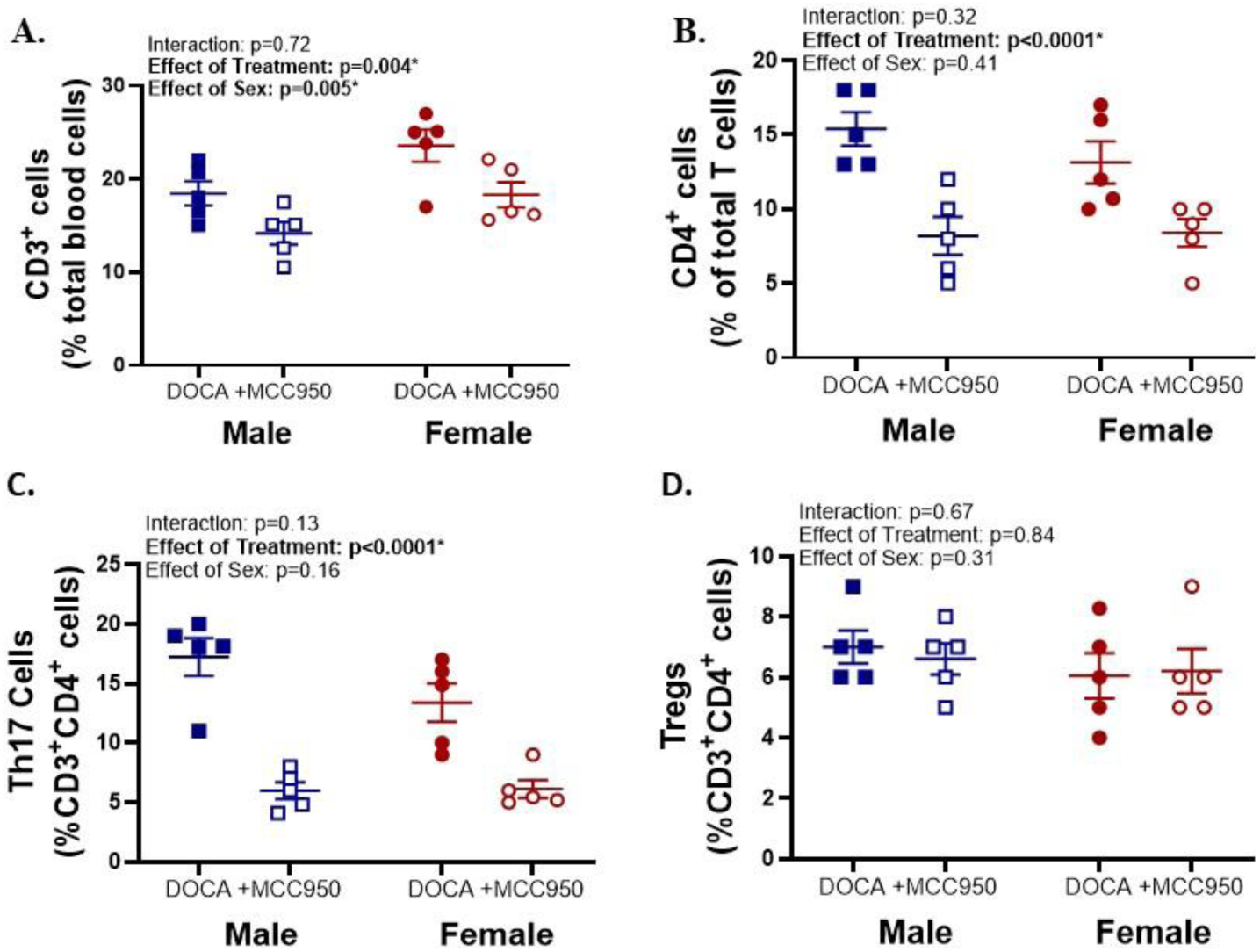
MCC950 reduces circulating pro-inflammatory T cells in DOCA-salt hypertensive males and females. T cell profiles were measured by flow cytometry in whole blood collected from deoxycorticosterone acetate (DOCA) treated male and female Sprague-Dawley (SD) rats randomized to saline control or the NLRP3 inflammasome inhibitor MCC950 (10 mg/kg/day in saline) for 3 weeks (n = 5). CD3^+^ T cells are expressed as a percentage of total blood cells (A), CD4^+^ T cells as a percentage of CD3^+^ T cells (B), T helper 17 (Th17) cells as a percentage of CD3^+^CD4^+^ T cells (C), and T regulatory cells (Tregs) as a percentage of CD3^+^CD4^+^ T cells (D). Data are expressed as mean ± SEM and analyzed using 2-way ANOVA. * indicates significance.

### MCC950 reduces renal pro-inflammatory T cells, with a more pronounced effect in males

The kidney plays a central role in BP control and renal inflammation has been linked to the development of hypertension ^17^. Therefore, renal T cells were also measured. The total percentage of CD3^+^ T cells in the kidney were not altered by MCC950 and were comparable between males and females (P_treatment_= 0.30, P_sex_= 0.81, P_interaction_= 0.82, Figure 4A). Consistent with our previous findings ^1^, males have more CD4^+^ T cells (P_sex_= 0.03) and Th17 cells (P_sex_= 0.001), while females have more anti-inflammatory Tregs (P_sex_= 0.02, Figure 4B-D). Inhibition of NLRP3 attenuated DOCA-induced increases in renal CD4+ T cells and Th17 cells, and this effect was greater in males (CD4^+^ T cells: P_treatment_= 0.0002, P_interaction_= 0.07; Th17 cells: P_treatment_< 0.0001, P_interaction_= 0.055). Tregs were not altered by MCC950 (P_treatment_= 0.78, P_interaction_= 0.78).

**Figure 4:**
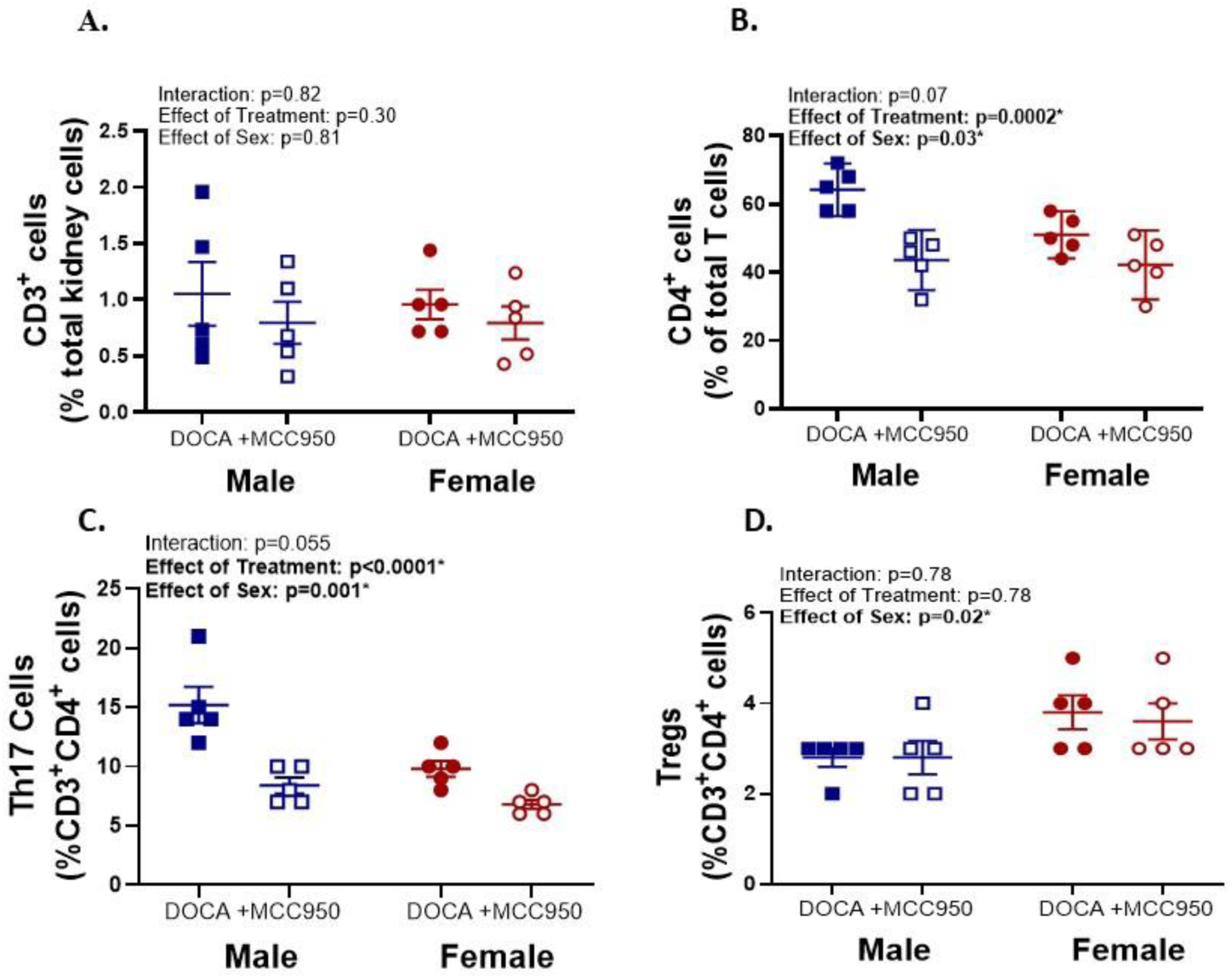
MCC950 lowers renal pro-inflammatory T cells in male and female DOCA-salt hypertensive rats with a more pronounced effect in males. T cell profiles were measured by flow cytometry in kidneys from deoxycorticosterone acetate (DOCA) treated male and female Sprague-Dawley (SD) rats randomized to saline control or the NLRP3 inflammasome inhibitor MCC950 (10 mg/kg/day in saline) for 3 weeks (n=5). CD3^+^ T cells are expressed as a percentage of total renal cells (A), CD4^+^ T cells as a percentage of CD3^+^ T cells (B), T helper 17 (Th17) cells as a percentage of CD3^+^CD4^+^ T cells (C), and T regulatory cells (Tregs) as a percentage of CD3^+^CD4^+^ T cells (D). Data are expressed as mean ± SEM and analyzed using 2-way ANOVA. * indicates significance.

### MCC950 reduces renal injury in both sexes of DOCA-salt hypertensive rats

Since both hypertension and inflammation contribute to renal injury ^18^, we evaluated the impact of NLRP3 inhibition on indices of renal injury in male and female DOCA-salt rats. Treatment with MCC950 attenuated DOCA-salt-induced increases in glomerular injury (PASH staining, P_treatment_= 0.0004, Figure 5A) and collagen deposition (Masson trichrome, P_treatment_= 0.046, Figure 5B) and the effects were comparable between males and females (P_sex_= 0.46, P_interation_= 0.71 and P_sex_= 0.47, P_interation_= 0.76, respectively). Despite this and consistent with the BP data, MCC950 improved creatinine clearance only in males (P_treatment_= 0.05; P_sex_= 0.003; P_interaction_=0.02, Figure S2A). Protein-to-creatinine ratio was not altered by MCC950 treatment (P_treatment_= 0.33; P_sex_= 0.33; P_interaction_= 0.35, Figure S2B).

**Figure 5:**
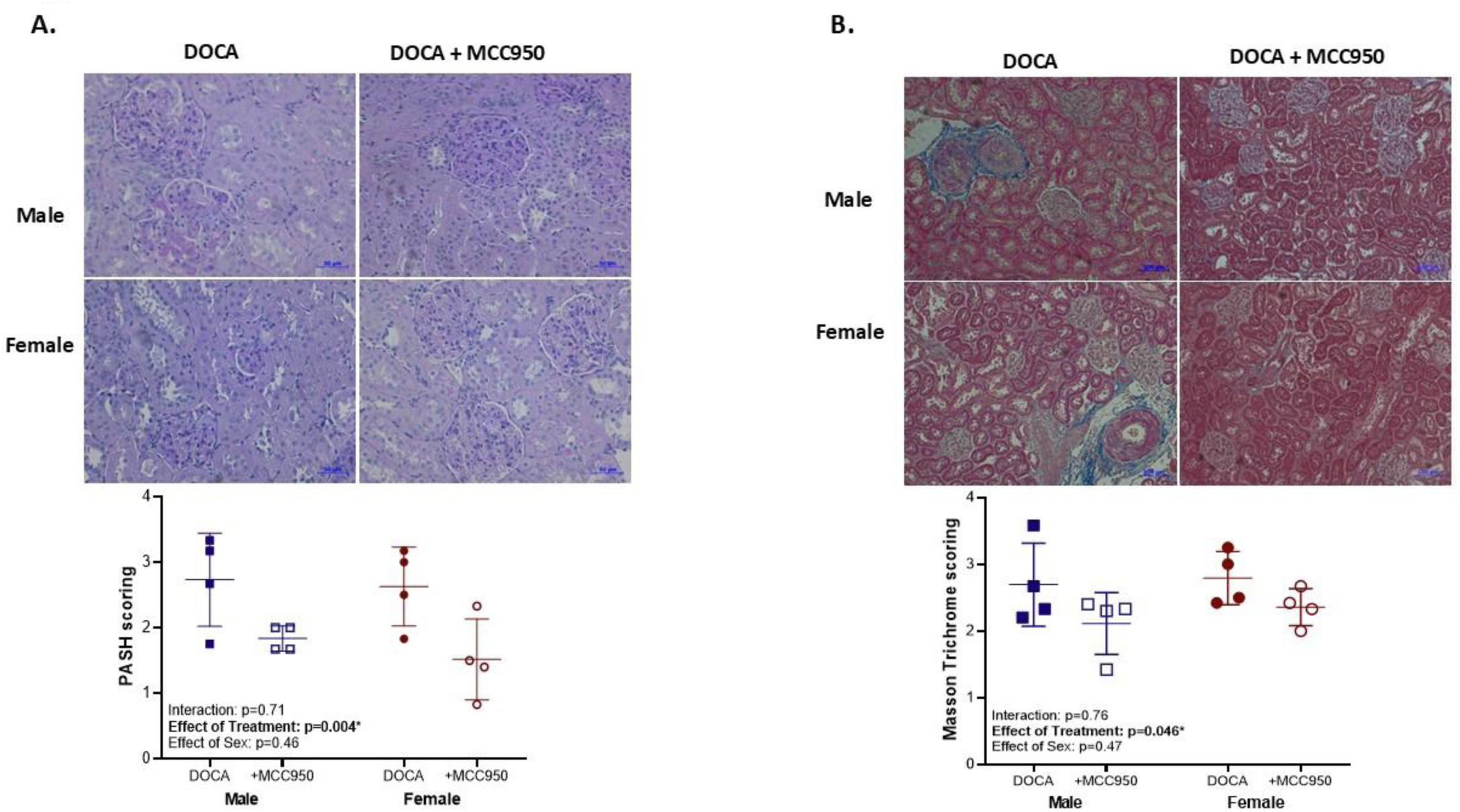
MCC950 reduces renal fibrosis and injury in male and female DOCA-salt hypertensive rats. Representative images at 200X magnification for PASH staining and scoring for glomerular injury (A), Masson trichrome staining and scoring for collagen deposition (B) in deoxycorticosterone acetate (DOCA) treated male and female Sprague-Dawley (SD) rats randomized to saline control or the NLRP3 inflammasome inhibitor MCC950 (10 mg/kg/day in saline) for 3 weeks (n = 4). Data are expressed as mean ± SEM and analyzed by 2-way ANOVA. * indicates significance.

## Discussion

The NLRP3 inflammasome has been implicated in the development of hypertension in male rodents ^9,10,19^. In this study, we extended this work to examine the NLRP3 inflammasome in normotensive and hypertensive males and females. Our findings show that although renal NLRP3 levels were similarly elevated in kidneys from hypertensive men and women, as well as in male and female DOCA-salt rats compared with normotensive controls, inhibition of NLRP3 with MCC950 significantly lowered BP only in male rats, despite comparable reductions in renal NLRP3 expression and T-cell infiltration in both sexes. These findings suggest a differential role for NLRP3 in BP regulation in males versus females.

Previous studies have shown that NLRP3 is activated in DOCA-salt hypertensive mice and in obese patients with hypertension ^9,20^. Consistent with these findings, renal NLRP3 levels were significantly greater in hypertensive patients and in DOCA-salt hypertensive rats vs. normotensive controls with no apparent sex differences. We further identified a sex-specific role for NLRP3 in the development of hypertension in male DOCA-salt rats. Pharmacological inhibition of NLRP3 with MCC950, a diarylsulfonylurea-containing compound shown *in vitro* and *in vivo* to selectively inhibit NLRP3 inflammasome activation, abolished the sex differences in BP ^10,21^. Once activated, the NLRP3 inflammasome leads to caspase-1 activation, which cleaves pro-IL-1β and pro-IL-18 into IL-1β and IL-18 propagating a pro-inflammatory response ^9,10^. Our findings confirmed the efficiency of MCC950 at the selected dose to reduce NLRP3 activation and its subsequent stimulation of caspase-1, IL-β and IL-18 in both sexes of DOCA-salt hypertensive rats.

Studies have shown that the NLRP3 inflammasome contributes to the development of renal inflammation and elevated BP in response to a variety of hypertensive stimuli including 1K/DOCA/salt and angiotensin II ^9,22,23^. Previous findings have shown that MCC950 treatment attenuates increases in BP in male 1K/DOCA/salt-induced hypertensive mice ^10^. In males, inflammasome deficient (ASC^-/-^) 1K/DOCA/salt mice also displayed blunted BP responses with decreased expression of renal inflammatory cytokines compared to wild-type (WT) 1K/DOCA/salt mice ^9^. In the current study, inhibition of NLRP3 with MCC950 only altered BP in males, despite similar NLRP3 expression in hypertensive males and females and similar decreases in NLRP3, IL-1β, IL-18 and caspase-1 with MCC950 in both sexes. The reason MCC950 treatment and decreases in NLRP3 did not alter BP in females remains unclear. However, IL-1β and caspase activity were lower in DOCA-salt females compared with males, and both pathways have been linked to the development of hypertension ^9,10^. Therefore, lower levels of these pro-hypertensive pathways may limit the impact of NLRP3 activation on BP in females. Regardless, our findings underscore the importance of sex-specific immune mechanisms in BP control.

T cells contribute to the incidence and progression of hypertension ^1,6,7,16^, and MCC950 treatment reduced renal T cells in male 1K/DOCA/salt mice ^10^. Consistent with this ^10^, MCC950 reduced pro-inflammatory CD4^+^ and Th17^+^ T cells in blood and kidneys of DOCA-salt hypertensive rats. The impact of MCC950 on pro-inflammatory immune cells was comparable in males and females, although males tended to have more pro-inflammatory T cells while females have more Tregs. Studies have shown that a subset of Tregs can suppress NLRP3 activation and alleviate monosodium urate-induced gouty arthritis ^24^. We have previously reported that greater Tregs in female rats attenuates DOCA-salt induced hypertension ^1^. NLRP3 inhibition selectively reduced renal Th17 cells without affecting Tregs, consistent with the established dependence of Th17 responses on inflammasome-mediated cytokine signaling ^25^. In contrast, Tregs seem less reliant on this pathway. Thus, we propose that greater Tregs in females may act as a counterbalance to mitigate the pro-hypertensive effects of NLRP3-driven inflammation ^1,3,7^ since renal Tregs remained higher in female DOCA-salt rats compared with males even after MCC950 treatment.

NLRP3 inhibition has also been linked to DOCA-salt hypertension-induced increases in renal inflammation and fibrosis ^9,10^. In the current study, we found that MCC950 treatment significantly improved creatinine clearance in male but not in female DOCA-salt rats, consistent with a BP lowering effect only in males. However, other makers of injury were comparable between males and females and reduced to a similar degree with NLRP3 inhibition. In addition, the protein to creatinine ratio was not altered by MCC950, suggesting that either the BP lowering effect in males was independent of the degree of renal injury or that the decrease in BP was insufficient to prevent injury. Indeed, even male rats treated with MCC950 remained hypertensive.

There are some limitations that should be considered when interpreting our findings. Only a single dose and duration of MCC950 were used, although MCC950 treatment has been shown to reduce renal inflammation, fibrosis, and injury even when administered 10 days after the establishment of 1K/DOCA/salt-induced hypertension ^10^. These data suggest that NLRP3 inhibition may not only prevent increases in BP but may also reverse established hypertension, which has important implications for therapeutic development. Our studies assessed the T cell profile, although other immune cells such as macrophages are also known to play a crucial role in BP control ^26^. Thus, future studies will explore the impact of MCC950 on other immune cells. In addition, the signal that activates the NLRP3 inflammasome in DOCA-salt hypertension was not examined. Studies have shown that increased osmolarity triggers NLRP3 oligomerization in cultured macrophages resulting in caspase-1 activation and IL-1β production ^27^. Treatment of isolated macrophages with aldosterone also stimulates caspase-1 activation and IL-18 production, and these changes were reduced with a mineralocorticoid receptor antagonist, suggesting that aldosterone could serve as a trigger ^28^. Clinically, mineralocorticoid receptor antagonists, including eplerenone and spironolactone, lower BP in patients with resistant hypertension ^29^. Thus, future studies will focus on determining if inflammasome inhibition contributes to the beneficial effects of mineralocorticoid receptor antagonists, and whether drugs that directly target the inflammasome, such as MCC950, would be effective therapies for resistant hypertension, especially in males.

## Non-standard Abbreviations and Acronyms

BP: Blood pressure
UNX: Uni-nephrectomy
DOCA: Deoxycorticosterone acetate
SD rats: Sprague Dawley rats
NLRP3: NOD-like receptor family pyrin domain containing 3
IL-1β: Interleukin-1 beta
IL-18: Interleukin-18
MCC950: Selective NLRP3 inflammasome inhibitor
Th17: T helper 17 cells
Tregs: Regulatory T cells
WT: Wild type

## Availability of data and materials

All raw data and materials are available upon request from the corresponding author.

## Author Contribution

MJR, EBT, AEL, JCS designed studies: KB, CML, KS and PH performed experiments, analyzed data prepared figures, and interpreted results of experiments; CML, AEL and JCS drafted manuscript; EBT, MJR and JCS edited and revised manuscript; DLM provided human kidney samples, KB, RM, EBT, RM, MJR, DLM and JCS approved final version of manuscript.

## Perspectives

Pharmacological inhibition of NLRP3 with MCC950 reduces BP only in male DOCA-salt rats abolishing sex difference in BP. In males, NLRP3 activation likely drives IL-1β-mediated T cell inflammation, contributing to elevated BP and renal injury ^10,30^. In females, BP regulation involves additional pathways such as greater Treg activity ^1^, which may reduce susceptibility to NLRP3-driven hypertension, inflammation and renal damage. As a result, targeting NLRP3 could be a promising therapeutic target in hypertension and its associated renal inflammation and injury in male patients ^31^, supporting the notion of using a personalized, sex-specific strategies in the treatment of hypertension.

## Novelty and Relevance

### What is New?

This study provides new evidence about the differential role for NLRP3 in BP regulation in male versus female DOCA-salt.

### What is Relevant?

There is increasing epidemiological evidence that men are more susceptible to hypertension and its associated renal injury than women. Pharmacological inhibition of the NLRP3 inflammasome has a sex specific effect to lower BP only in male DOCA-salt hypertensive rats, despite similar reduction in T cell infiltration and renal injury in both sexes.

### Clinical/Pathophysiological Implications

Targeting NLRP3 could be a promising therapeutic target in hypertension and its associated renal inflammation and injury and could act as an adjunct treatment to improve efficiency of mineralocorticoid receptor antagonists for patients with resistant hypertension.

